# Phosphorylation of the mitochondrial triphosphate tunnel metalloenzyme TTM1 regulates programmed cell death in senescence

**DOI:** 10.1101/2020.10.06.328278

**Authors:** Purva Karia, Keiko Yoshioka, Wolfgang Moeder

## Abstract

The role of mitochondria in programmed cell death (PCD) during animal growth and development is well documented, but much less is known for plants. We previously showed that the *Arabidopsis thaliana* triphosphate tunnel metalloenzyme (TTM) proteins TTM1 and TTM2 are tail-anchored proteins that localize in the mitochondrial outer membrane and participate in PCD during senescence and immunity, respectively. Here, we show that TTM1 is specifically involved in senescence induced by abscisic acid (ABA). Moreover, phosphorylation of TTM1 by multiple mitogen-activated protein kinases (MAPKs) regulates its function and turnover. A combination of proteomics and *in vitro* kinase assays revealed three major phosphorylation sites of TTM1 (S10, S437, and S490), which are phosphorylated upon perception of senescence cues such as ABA and prolonged darkness. S437 is phosphorylated by the MAP kinases MPK3 and MPK4, and S437 phosphorylation is essential for TTM1 function in senescence. These MPKs, together with three additional MAP kinases (MPK1, MPK7, and MPK6), phosphorylate S10 and S490, marking TTM1 for protein turnover, which likely prevents uncontrolled cell death. Taken together, our results show that multiple MPKs regulate the function and turnover of the mitochondrial protein TTM1 during senescence-related PCD, revealing a novel link between mitochondria and PCD.

**Summary:** Email addresses

## INTRODUCTION

Senescence is a type of highly regulated developmental programmed cell death (PCD) in which nucleic acids, carbohydrates, lipids, and proteins are remobilized from senescing tissues such as leaves to other plant tissues such as seeds or roots. Senescence is crucial for ending the life cycle of plants and for survival under unfavorable conditions (Daneva et al., 2016). During senescence, chloroplasts are the first organelles to be degraded and leaf yellowing resulting from the loss of chlorophyll provides a visible sign of senescence (Keech et al., 2007; Wada et al., 2009). Mitochondria play crucial roles in senescence, as a signaling hub and by providing energy to the cell while chloroplasts are disassembled (Kmiecik et al., 2016; Chrobok et al., 2016). However, the precise molecular mechanism by which mitochondria contribute to plant PCD remains elusive.

In addition to being triggered by developmental cues, senescence can be triggered by abiotic stresses such as drought or prolonged darkness. Exogenous application of phytohormones that are related to these external stresses, such as abscisic acid (ABA), ethylene, or jasmonic acid (JA), also trigger senescence (Weaver et al., 1998, Kim et al., 2011; Liu et al., 2016). It has long been known that ABA levels increase in senescing leaves (Samet and Sinclair, 1980). However, although ABA signal transduction and the role of ABA in seed germination, stomata closure, and drought have been studied extensively (Cutler et al., 2010; Sussmilch et al., 2019; Yan and Chen, 2017), relatively little is known about the role of ABA in leaf senescence. The senescence-related transcription factor ARABIDOPSIS NAC DOMAIN CONTAINING PROTEIN (NAP) induces the expression of *ABSCISIC ALDEHYDE OXIDASE3* (*AAO3*), whose protein product catalyzes the final step in ABA biosynthesis. ABA then promotes the transcription of chlorophyll catabolic genes such as *NON-YELLOW COLORING1* (*NYC1*), and *PHEOPHORBIDE a OXYGENASE* (*PAO*) via the ABA-RESPONSIVE ELEMENT (ABRE) BINDING PROTEINS ABF2, ABF3, and ABF4 transcription factors (Yang et al., 2014, Gao et al., 2016).

ABA perception has been elucidated in detail: ABA binds to the PYRABACTIN RESISTANCE1/PYR1-LIKE/REGULATORY COMPONENTS OF ABA RECEPTOR (PYR1/PYL/RCAR) ABA receptors, leading to the inactivation of Clade A type 2C protein phosphatases (PP2Cs), which then release the inhibition of the SUCROSE NON-FERMENTING-1 RELATED PROTEIN KINASE 2 (SnRK2) kinases (Cutler et al., 2010). Activated SnRK2 kinases (SnRK2.2, SnRK2.3, and SnRK2.6) then phosphorylate downstream targets such as ABF transcription factors, which in turn induce the expression of many ABA-responsive genes (Fujita et al., 2009; Fujii et al., 2009) and other target genes (Wang et al., 2013; Umezawa et al., 2013). The *snrk2*.*2 snrk2*.*3 snrk2*.*6* (*snrk2*.*2/3/6*) triple mutant is insensitive to ABA (Fujii et al., 2009) and displays delayed ABA-induced senescence (Gao et al., 2016; Zhao et al., 2016). Previous work suggested that these SnRK2 kinases activate the MAP kinase kinase kinases MAPKKK18 and MAPKKK17, which in turn activate the MAP kinase kinase MKK3 (Danquah et al., 2015; Tajdel et al., 2016; Matsuoka et al., 2018). MKK3, as well as ABA treatment, activate the group C MAP kinases MPK1, MPK2, MPK7, and MPK14. The *mapkkk17 mapkkk18* double mutant and the *mkk3* mutant exhibit hypersensitivity to ABA treatment during germination and altered expression of ABA-inducible genes (Danquah et al., 2015).

In our previous study, we showed that triphosphate tunnel metalloenzyme 1 (TTM1) plays a positive role in the regulation of senescence, as *ttm1* mutant plants displayed delayed natural and dark-induced senescence (Ung and Karia et al., 2017). TTM proteins are characterized by a unique tunnel structure, composed of eight antiparallel β-strands forming a β-barrel, and a characteristic EXEXK motif (where X is any amino acid) that is crucial for their enzymatic activity (Lima *et al*., 1999; Iyer and Aravind, 2002; Gallagher et al., 2006). Many TTM proteins also possess additional domains, such as a nucleotide kinase (P-loop kinase) or a conserved histidine α-helical (CHAD) domain (Iyer and Aravind, 2002), which also suggests that these TTM proteins act on nucleotide or polyphosphate substrates (Lorenzo-Orts et al., 2019).

To date, an enzymatic function has only been reported for a few TTM proteins, which have diverse activities including RNA triphosphatase, adenylate cyclase, thiamine triphosphatase, and tripolyphosphatase (Iyer and Aravind; 2002; Bettendorff and Wins, 2013, Moeder et al., 2013). Known TTM substrates include nucleotides and organophosphates such as ATP, thiamine triphosphate, the 5′ end of nascent RNAs, and tripolyphosphate (Lima et al., 1999; Gallagher et al., 2006; Lakaye et al., 2004; Keppetipola et al., 2007; Moeder et al., 2013). In addition, TTMs require a divalent metal cation cofactor, usually magnesium (Mg^2+^) or manganese (Mn^2+^) (Bettendorff and Wins, 2013).

Plant genomes usually encode two types of TTM proteins: a small TTM protein containing only one TTM domain, termed TTM3 in Arabidopsis (*Arabidopsis thaliana*) (Moeder et al., 2013; Martinez et al., 2015), and another type having a fusion of a TTM and a P-loop kinase domain (Iyer and Aravind, 2002; Ung et al., 2014; Ung and Karia et al., 2017). Arabidopsis has two TTM proteins of the second type, TTM1 and TTM*2*, each with a P-loop kinase/uridine kinase-like domain (UK) followed by a TTM domain (Ung et al., 2014). Both proteins also have an extended C-terminal tail region containing a coiled-coil domain and a transmembrane (TM) domain. Previously, we reported that Arabidopsis TTM1 and TTM2 are tail-anchored proteins that are localized to the mitochondrial outer membrane via their TM domain (Ung and Karia et al., 2017; Kriechbaumer et al., 2009). Most dicot plants have both TTM1 and TTM2 homologues, while monocots have only one homologue (in addition to one TTM3 homologue). We demonstrated that both TTM1 and TTM2 display pyrophosphatase activity (Ung et al., 2014; Ung and Karia et al., 2017), which is unique among TTM proteins, as they usually hydrolyze triphosphate substrates (Bettendorff and Wins, 2013).

Although TTM1 and TTM2 share over 92% sequence similarity and 66% sequence identity, which is even higher in the UK (81% identity) and TTM domains (73% identity), they play distinct roles in plant immunity (Ung et al., 2014) and senescence (Ung and Karia et al., 2017). Their biological roles are determined by their expression patterns, since *TTM2* can complement the *ttm1* mutant phenotype when expressed under the control of the *TTM1* promoter, and *vice versa* (Ung and Karia et al., 2017). Each protein is also connected to a different type of PCD: TTM2 negatively regulates immunity-related cell death (Ung et al., 2014), whereas TTM1 positively regulates senescence-associated cell death (Ung and Karia et al., 2017).

Here, we show that the mitochondrial outer membrane protein TTM1 is regulated by senescence cues such as dark or ABA through its phosphorylation and accelerates senescence-related PCD. TTM1 is phosphorylated at multiple serine residues upon treatment with ABA, and phosphorylation at serine 437 (S437) is required for its function. We identified multiple MAP kinases that phosphorylate TTM1 at S437, as well as at two other residues, S10 and S490. Our data indicate that phosphorylation at these two additional positions marks TTM1 for degradation by the 26S proteasome, revealing a role for TTM1 in linking senescence, ABA, and mitochondria.

## RESULTS

### TTM1 is involved in ABA-induced senescence

In our previous study, we showed that TTM1 plays a positive regulatory role in senescence, as *ttm1* mutant plants displayed delayed natural and dark-induced senescence (Ung and Karia et al., 2017). The phytohormones ABA, ethylene, and JA are associated with senescence (Breeze et al., 2011), and treatment with any of these hormones leads to senescence/chlorophyll degradation over the course of 3–4 days (Kim et al., 2011). Therefore, we tested whether TTM1 is a general senescence regulator or is part of a specific phytohormone-associated senescence pathway.

To this end, we quantified chlorophyll levels from detached leaves after floating them on water containing ABA, the ethylene precursor 1-aminocyclopropane-1-carboxylic acid (ACC), or methyl jasmonate (MeJA). MeJA and ACC treatment induced accelerated senescence, as seen by reduced chlorophyll levels in wild-type Columbia (Col) plants, compared to buffer control (Figure 1A). By contrast, we observed no differences between treated *ttm1* and wild-type plants (Figure 1A). However, we noticed a substantial delay in chlorophyll degradation in ABA-treated *ttm1* leaves (Figure 1B). The delay in *ttm1* was comparable to that observed in the ABA-insensitive mutant *abi5* (Figure 1B), indicating that TTM1 is part of the ABA-mediated senescence signaling pathway.

**Figure 1.**
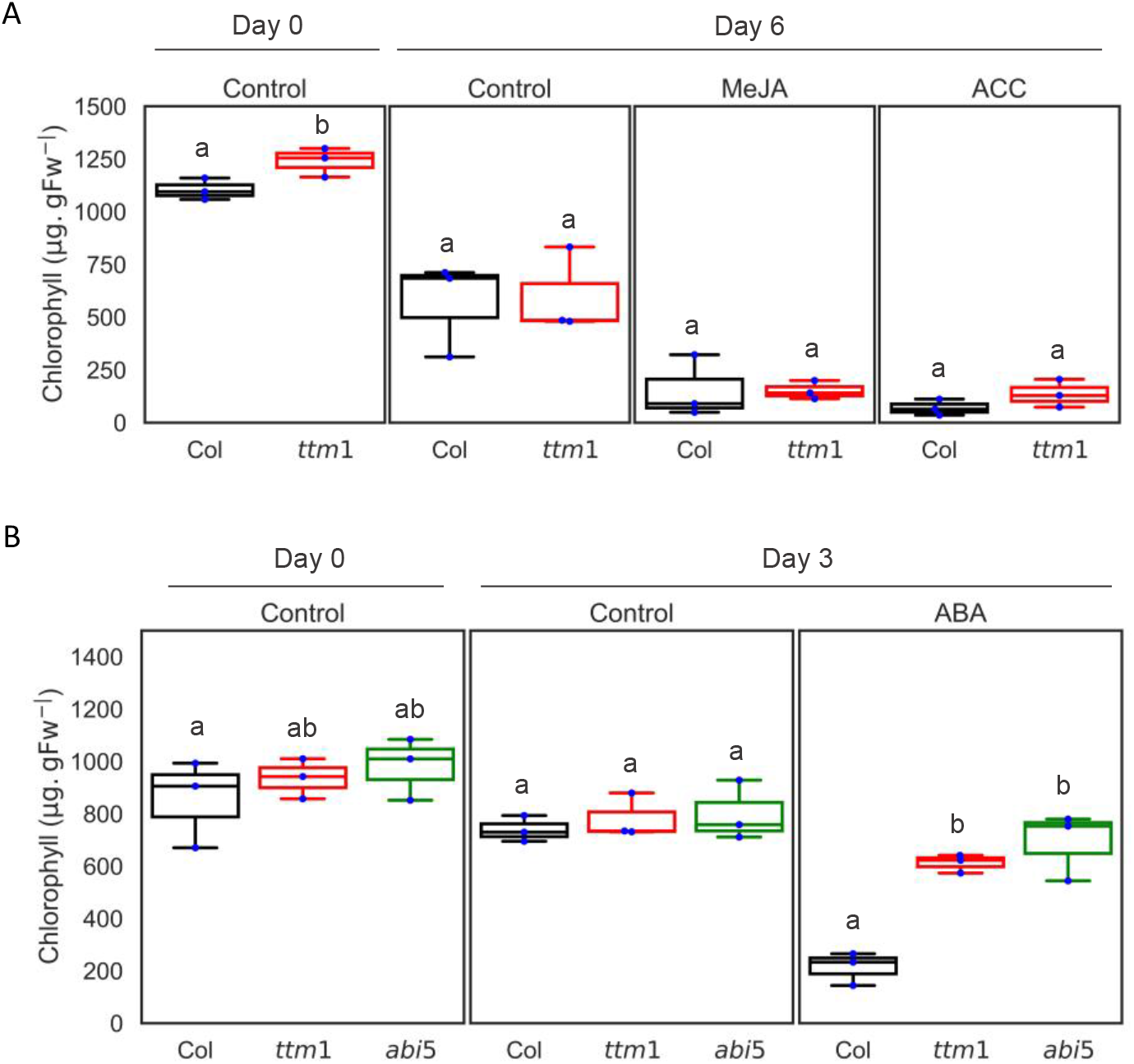
*ttm1* displays delayed leaf senescence upon ABA treatment. **(A)**. Leaves of 4-5-week-old *Arabidopsis thaliana* accession Columbia (Col) wild type and *ttm1* mutants were detached and floated for 6 days on de-ionized water ± 25 µM MeJA or de-ionized water containing 3 mM MES (pH 5.8) ± 50 µM ACC (Controls: ACC, 3 mM MES, MeJA, 0.1% ethanol). **(B)** 4-5-week-old *Arabidopsis thaliana* wild type (Col), *ttm1, aao3* and *abi5* mutant leaves were detached and floated for 3 days on de-ionized water +/- 50 µM ABA (Control, 0.1% ethanol). *abi5* served as a positive (ABA insensitive) control. Leaves were under 16/8 hr light/dark conditions. Total chlorophyll content was measured from samples before and after treatment. Each bar represents the mean ± SE (n = 3). ABA and MeJA experiments were repeated three times and the ACC experiment was performed twice. Measurements are plotted as box plots displaying the first and third quartiles, split by the median; whiskers extend to a maximum of 1.5× interquartile range beyond the box. Statistical analysis was performed using (A) student’s t-test and (B) one-way ANOVA with Tukey’s HSD post hoc test. Different letters denotes a significant difference between two samples.

### TTM1 is phosphorylated upon dark treatment

*TTM1* is transcriptionally up-regulated during dark-induced senescence (Ung and Karia et al., 2017). Therefore, we tested whether *TTM1* is transcriptionally regulated by ABA. At 24 h, ABA treatment induced the expression of the ABA marker gene *RESPONSIVE TO DESICCATION 29A* (*RD29A*) (Supplemental Figure S1). At 72 h, ABA treatment caused the up-regulation of the senescence marker gene *SENESCENCE-ASSOCIATED GENE 12* (*SAG12*), coinciding with chlorophyll loss. By contrast, we observed no transcriptional up-regulation of *TTM1* after treatment with ABA (Supplemental Figure S1A). This observation is also supported by publicly available microarray datasets (Supplemental Figure S1B; Goda et al., 2008).

These data indicated that ABA does not transcriptionally regulate *TTM1* expression. Therefore, we hypothesized that TTM1 might be regulated post-translationally. Indeed, several proteomics studies have reported increased phosphorylation of TTM1 at S437 upon ABA treatment (Kline et al., 2010; Wang et al., 2013). These observations prompted us to investigate the phosphorylation status of TTM1.

First, we tested whether an increase in S437 phosphorylation occurred during dark treatment. For this purpose, we transiently overexpressed yellow fluorescent protein (YFP)-tagged TTM1 in *Nicotiana benthamiana* leaves. This system has successfully been used to induce senescence/cell death (Ung and Karia et al., 2017). *YFP-TTM1* has been shown to functionally complement the delayed senescence phenotype of *ttm1* similar to untagged *TTM1* (Ung and Karia et al., 2017).

To measure phosphorylation, YFP-TTM1 was immunoprecipitated from dark-treated and control *N. benthamiana* leaves transiently expressing the *35Spro:YFP-TTM1* construct. Our LC-MS/MS analysis after immunoprecipitation and trypsin digestion identified 172 and 262 total spectra for TTM1 in control and dark-treated samples, respectively, with a minimum of 2 peptides and a 0.8% false discovery rate (FDR). We observed a 50% increase in phosphorylation at S437 in dark-treated samples relative to controls (Supplemental Table 1), showing that S437 is phosphorylated both during dark-induced senescence and after ABA treatment (Kline et al., 2010; Wang et al., 2013).

### Phosphorylation at S437 is required for TTM1 function

Considering that ABA and dark treatment both lead to TTM1 phosphorylation at S437, we hypothesized that phosphorylation at this position is required for TTM1 function. To test this idea, we expressed phospho-dead (*TTM1*^*S437A*^) and phospho-mimetic (*TTM1*^*S437D*^) variants of *TTM1* under the control of the native *TTM1* promoter, in *ttm1* knockout plants and performed functional complementation assays (Ung and Karia et al., 2017). We confirmed that all complementation lines had comparable expression levels for all *TTM1* variants by RT-PCR (Supplemental Figure S2).

As expected, *ttm1* mutant plants showed a significant delay in senescence, as they retained significantly more chlorophyll than wild type at day 3 after ABA treatment (Figure 2A, B). Also as expected, two independent transgenic *ttm1* lines expressing wild-type *TTM1* (*TTM1pro:TTM1*) complemented the senescence phenotype back to that seen in wild-type (Figure 2A, B, Ung and Karia et al., 2017). Similarly, the phospho-mimetic *TTM1pro:TTM1*^*S437D*^ construct complemented the *ttm1* phenotype. However, the construct for the phospho-dead variant of *TTM1, TTM1pro:TTM1*^*S437A*^, did not complement the *ttm1* mutant (Figure 2A, B).

**Figure 2:**
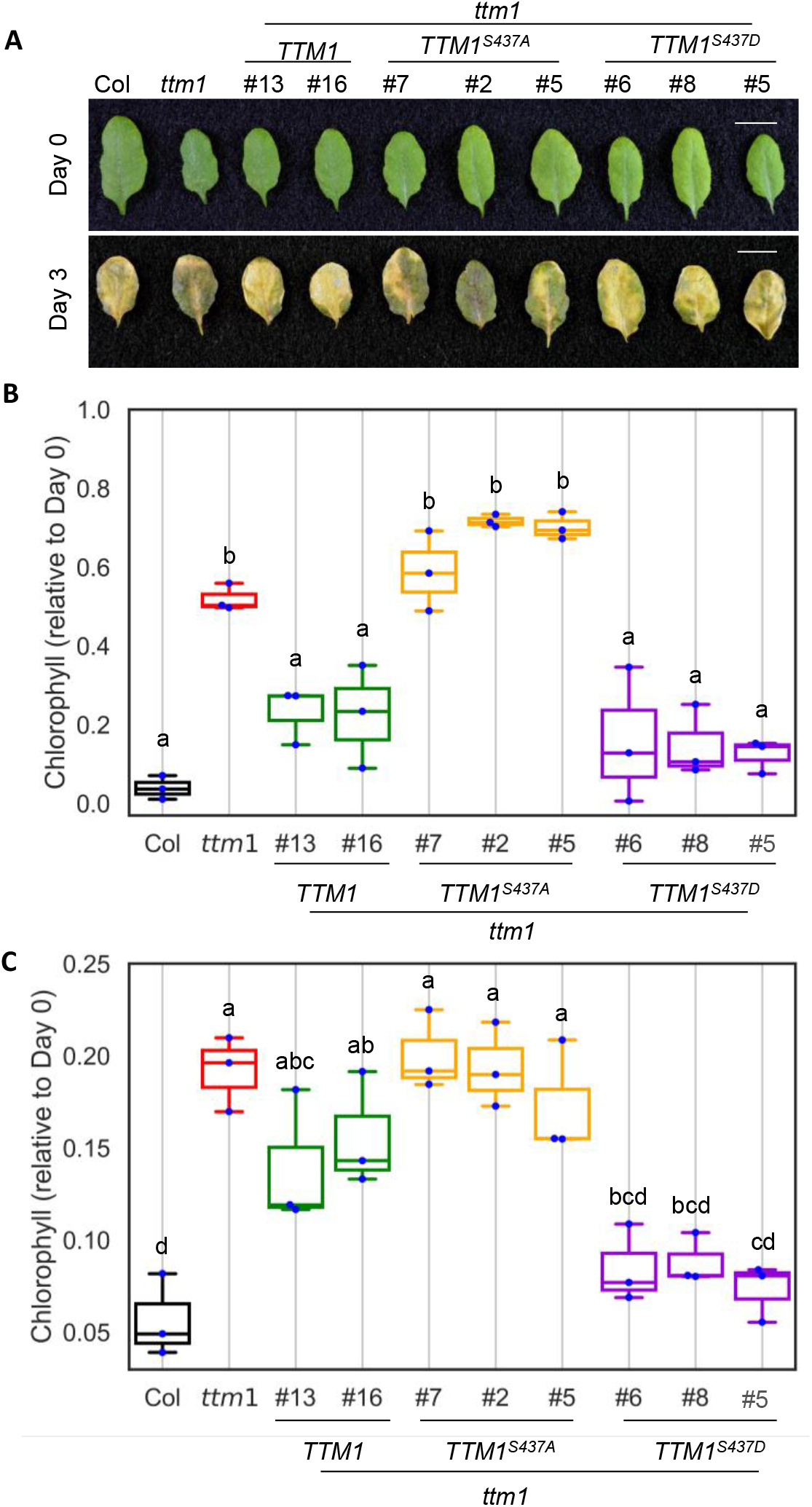
TTM1^S437A^ does not complement the delayed senescence phenotype of *ttm1*. **(A, B)** Leaves of 4-5-week-old *Arabidopsis thaliana* wild type (Col), *ttm1*, two independent *ttm1/TTM1* lines, three independent *ttm1/TTM1*^*S437A*^ and three independent lines *ttm1/TTM1*^*S437D*^ were floated for 3 days on water + 50 µM ABA. **(A)** Images of leaves on day 0 and 3 after ABA treatment. **(B)** Total chlorophyll content was measured from samples before/after treatment, normalized values of chlorophyll content after treatment are plotted. **(C)** Leaves from the same lines were detached and floated for 5 days on water in the dark and chlorophyll was quantified. Both experiments were repeated independently three times. Measurements are plotted as box plots displaying the first and third quartiles, split by the median; whiskers extend to a maximum of 1.5× interquartile range beyond the box. Statistical analysis was performed using a one-way ANOVA with Tukey’s HSD post hoc test. Different letters denote a significant difference between samples.

We tested the same lines in the dark-induced senescence assay, which confirmed that only the phospho-dead version of *TTM1* failed to complement the *ttm1* mutant phenotype (Figure 2C), indicating that phosphorylation at S437 upon ABA and dark treatment is required for TTM1 function. Taken together, these data indicate that phosphorylation at S437 is essential for TTM1 function to induce senescence-associated cell death.

### TTM1 is targeted by multiple MAP kinase

In order to understand the regulation of TTM1 and its role in senescence signaling, we aimed to identify the kinase(s) that phosphorylate TTM1 upon perception of senescence cues such as ABA and dark treatment. The ABA-induced phosphorylation of TTM1 was greatly reduced in the *snrk2*.*2/2*.*3/2*.*6* triple mutant (Wang et al., 2013). Thus, we first hypothesized that these SnRK2 kinases might phosphorylate TTM1 and tested this by an *in vitro* kinase assay using purified glutathione S-transferase (GST)-tagged SnRK2.2, SnRK2.3 and SnRK2.6 protein and maltose binding protein (MBP)-His-tagged TTM1. However, we detected only the auto-phosphorylation of SnRK2.6 protein on the autoradiograph, and no phosphorylation of the recombinant MBP-His-TTM1 protein (Supplemental Figure S3A).

The fact that S437 is followed by a proline residue suggested that S437 might be phosphorylated by a MAP kinase (Sörensson et al., 2012; Rayapuram et al., 2018). The *Arabidopsis thaliana* genome encodes 20 MAP kinases (Xu and Zhang, 2015). The group C kinases MPK1, MPK2, MPK7, and MPK14 had previously been shown to be activated by ABA (Danquah et al., 2015). MPK12 is also activated by ABA (Jammes et al., 2009). MPK6 from group A has also been connected to senescence (Zhou et al., 2009). In addition, publicly available transcriptome datasets indicate that *MPK3, MPK4, MPK11*, and *MPK5* are transcriptionally up-regulated in senescing leaves (Klepikova et al., 2016; Supplemental Figure S4). We therefore tested these ten MAP kinases with connections to ABA or senescence through *in vitro* kinase assays. Of the four group C kinases, MPK1 and MPK7, but not MPK2 or MPK14, phosphorylated TTM1 (Figure 3A). MPK6 also phosphorylated TTM1 *in vitro* (Figure 3A), as did MPK3 and MPK4 (Figure 3B), but not MPK5, MPK11, or MPK12 (Supplemental Figure S3B).

**Figure 3:**
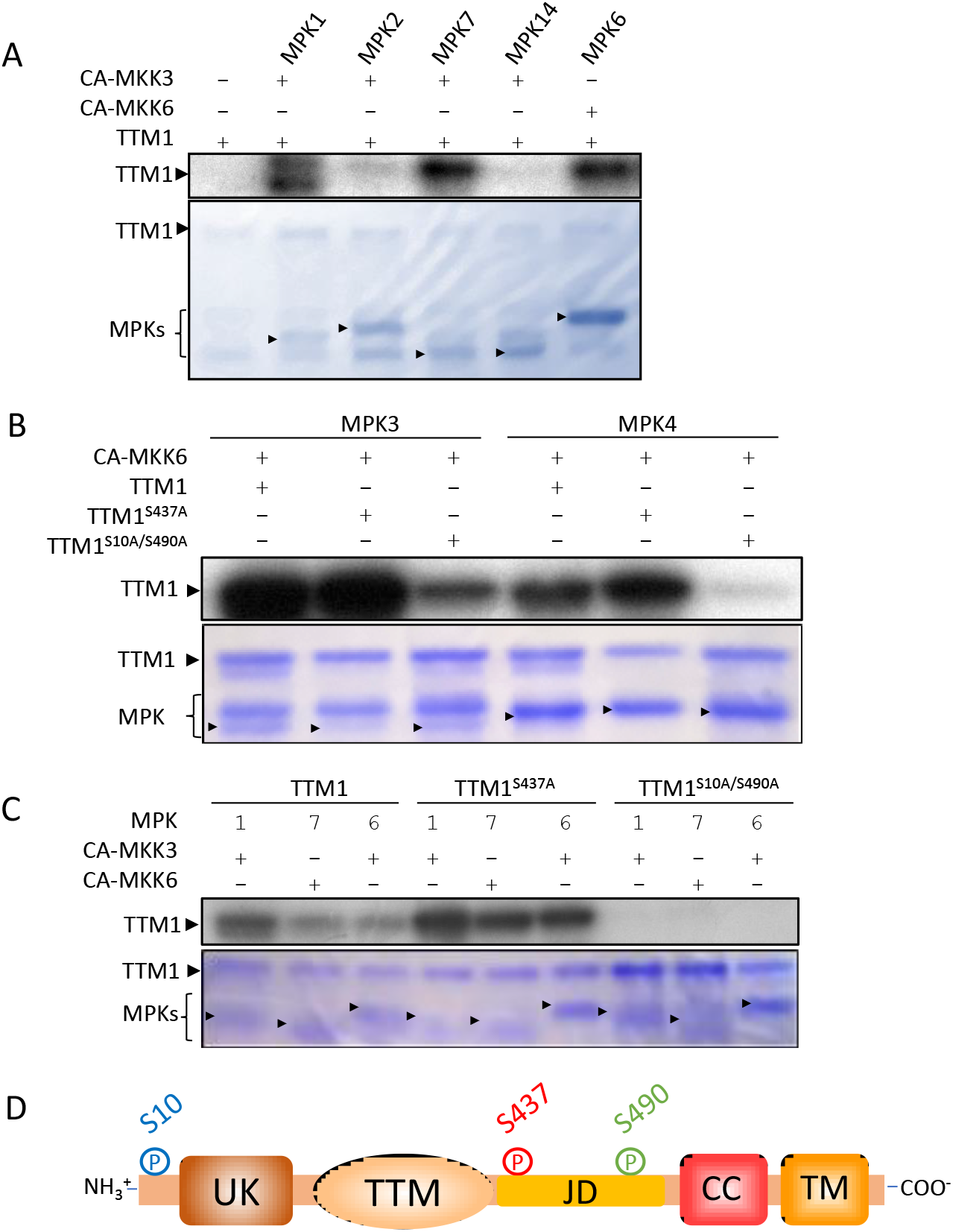
*In vitro* phosphorylation of TTM1 by MAP kinases. **(A-C)** Autoradiographs (top panel) and Coomassie R-250 stained SDS-PAGE (bottom panel) are shown. All MPKs used were GST-tagged and TTM1 variants were MBP-His tagged. The presence of proteins in each of the reactions is shown above the images. The MPK and TTM bands are marked by arrows. **(A)** Phosphorylation of TTM1 by different MAP kinases. **(B)** Phosphorylation of TTM1, TTM^S10A/S490A^, and TTM1^S437A^ by MPK1, MPK7, and MPK6. **(C)** Phosphorylation of TTM1, TTM^S10A/S490A^, and TTM1^S437A^ by MPK3 and MPK4 **(D)** Positions of the phosphorylation sites relative to protein domains. UK = Uridine kinase domain, TTM =TTM domain, JD = Junction domain, CC = Coiled-coil domain, TM = Transmembrane domain

To test if these MAP kinases phosphorylate TTM1 at the functionally essential S437 residue, we next tested the phospho-dead variant TTM1^S437A^ for phosphorylation. However, TTM1^S437A^ still acted as a substrate for MPK1, MPK6, and MPK7 in our *in vitro* kinase assay (Figure 3C). This data suggested that these MAP kinases may phosphorylate residues in TTM1 other than S437. Thus, to identify the other phosphorylation sites, we conducted an LC-MS/MS phospho-peptide analysis using MPK7 as kinase. MPK7 did not phosphorylate TTM1 at S437 as predicted, but did phosphorylate two other residues: S10, located at the N-terminus before the uridine kinase domain, and S490 in the junction domain before the coiled-coil domain (Table 1; Figure 3D). We thus conducted *in vitro* phosphorylation assay with the kinases MPK1, MPK6, and MPK7, using a TTM1^S10A/S490A^ double phospho-dead mutant. As expected, we detected no phosphorylation of TTM1^S10A/S490A^ by MPK1, MPK6, or MPK7 (Figure 3C), demonstrating that these three kinases phosphorylate S10 and S490 rather than S437.

This finding prompted us to expand our analysis of TTM1 phosphorylation with a LC-MS/MS phospho-peptide analysis of MPK3 and MPK4 *in vitro* phosphorylation products. Interestingly, both kinases phosphorylated not only S437, but also S10 and S490, thereby adding MPK3 and MPK4 to the list of kinases targeting these two residues (Table 1). The *in vitro* kinase assay showed no reduction of phosphorylation of TTM1^S437A^ by either kinase, but reduced levels of TTM1^S10A/490A^ phosphorylation by MPK3 and almost no TTM1^S10A/490A^ phosphorylation by MPK4, confirming the phospho-peptide analysis (Figure 3B).

Taken together, these results imply that MPK3 and MPK4 regulate TTM1 function via phosphorylation at S437. Furthermore, we identified additional kinases, including MPK1, MPK6, MPK7, MPK3, and MPK4, that phosphorylate TTM1 at two additional sites, S10 and S490.

### Phosphorylation status of TTM1 regulates its function and turnover

Transient expression of wild-type *TTM1* in *N. benthamiana* leaves causes cell death when plants are maintained in the dark for prolonged time suggesting that TTM1 functions as a positive regulator of dark-induced cell death (Ung and Karia et al., 2017). To further investigate the role of TTM1 phosphorylation at S10 and S490, we tested the effect of transient overexpression of wild-type, phospho-mimetic, and phospho-dead variants of TTM1 on dark-induced cell death in *N. benthamiana* leaves. The single (S437A), double (S10A, S490A), and triple phospho-dead TTM1 variants displayed a similar degree of cell death to wild-type TTM1, indicating that the phosphorylation of these residues does not affect protein function itself (Figure 4A). By contrast, the TTM1 ^S437D^ single variant displayed cell death similar to TTM1 wild type, whereas both the TTM1 ^S10D/S490D^ double and TTM1 ^S10D/S437DS490D^ triple variants showed a significant reduction in the cell death phenotype where the phospho-mimetic triple variant exhibited almost 50% reduction in cell death (Figure 4B, C). These results suggest that unlike S437, phosphorylation at S10 and S490 negatively affects the ability of TTM1 to induce cell death.

**Figure 4:**
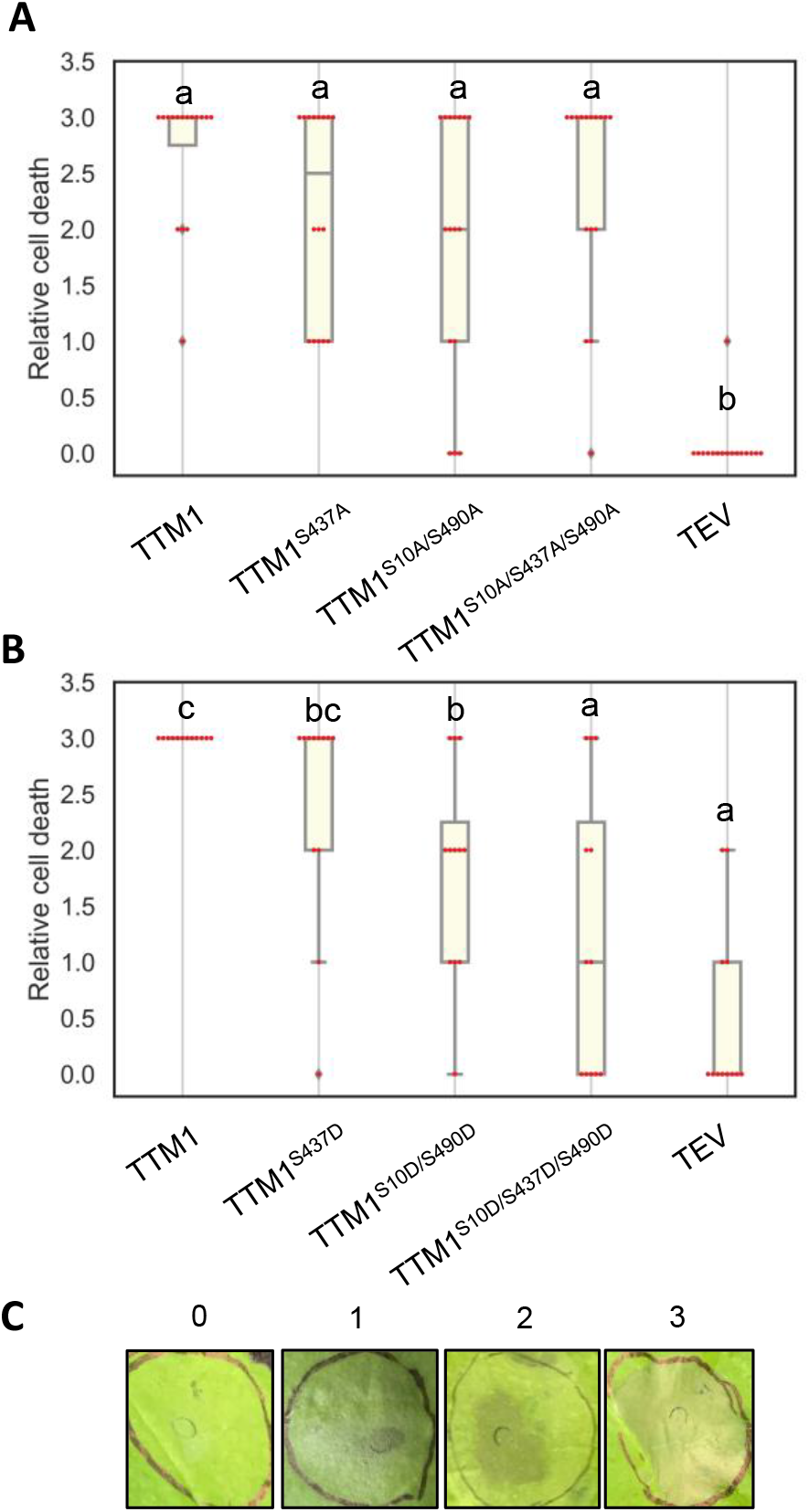
Double and triple phospho-mimetic TTM1 variants show reduced cell death upon dark treatment. **(A, B)** Cell death caused by the expression of TTM1 variants. *N. benthamiana* plants expressing YFP-tagged TTM1 and **(A)** phospho-dead variants (TTM1^S437A^, TTM1^S10A/S490A^, TTM1^S10A/S437A/S490A^), and **(B)** phospho-mimetic variants (TTM1^S437D^, TTM1^S10D/S490D^, TTM1^S10D/S437D/S490D^) were shifted to the dark 24 hpi. Agrobacterium carrying HcPro from tobacco etch virus (TEV) was coninfiltrated to suppress gene silencing and TEV alone served as a negative control. Cell death was scored after 9 days in the dark. **(C)** Examples of cell death areas. 3: Strong cell death, 2: mild cell death, 1: weak cell death, 0: No cell death. Each bar represents the mean ± SE (n >= 12), the experiment was repeated 3 times with similar results.

TTM1 is a tail-anchored protein that localizes to the mitochondrial outer membrane (Ung and Karia et al., 2017). We analyzed the protein levels and localization of the different phosphorylation mutants by confocal microscopy. As reported previously, wild-type TTM1 showed a punctate pattern (Figure 5A; Ung and Karia et al., 2017). A similar punctate pattern was also observed with the phospho-dead and phospho-mimetic single variants at the S437 residue (TTM1^S437D^, TTM1^S437A^), indicating that the phosphorylation status at S437 does not affect subcellular localization or protein degradation (Figure 5B, E). However, we noticed a strong reduction in puncta and the appearance of a cytosolic pattern for the phospho-mimetic variants TTM1^S10D/S490D^ and TTM1^S10D/S437D/S490D^ (Figure 5F, G), while the corresponding phospho-dead variants displayed a normal localization pattern (Fig 5C, D).

**Figure 5.**
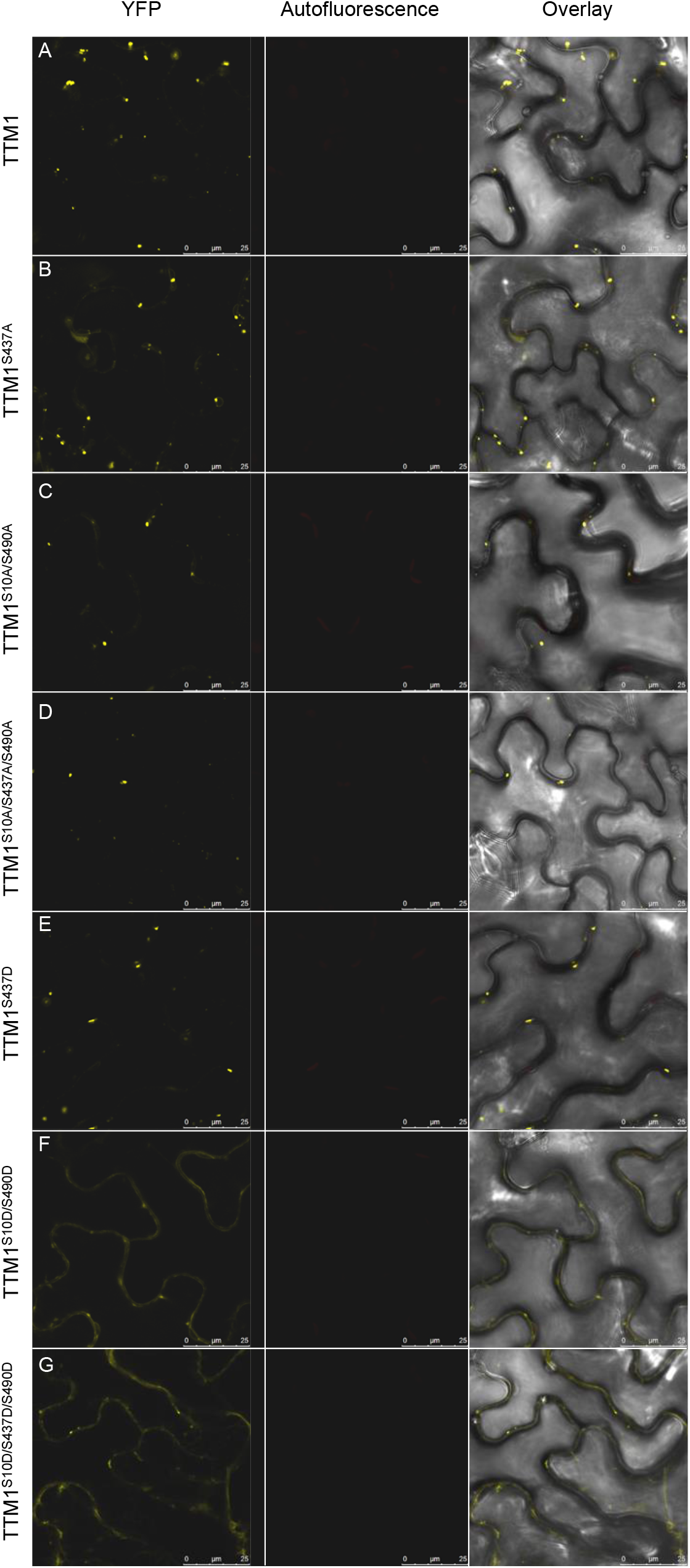
Double and triple phospho-mimetic TTM1 variants show alteration in their subcellular localization. Subcellular localization of TTM1 variants**: (A)** TTM1 wild type, phospho-dead variants **(B)** TTM1^S437A^, **(C)** TTM1^S10A/S490A^ and **(D)** TTM1^S10A/S437A/S490A^, and the phospho-mimetic variants **(E)** TTM1^S437D^, **(F)** TTM1^S10D/S490D^, and **(G)** TTM1^S10D/S437D/S490D^. All YFP-tagged variants were transiently overexpressed in *N. benthamiana*. Leaf discs were visualized by confocal microscopy at 48 hpi. Left panel: YFP signal, Middle panel: Auto fluorescence, Right panel: Overlay.

This raised the possibility that phosphorylation at S10 and S490 may lead to enhanced turnover of the TTM1 protein, possibly via the 26S proteasome. Therefore, we monitored YFP-TTM1 protein levels in the presence and absence of the proteasome inhibitor MG132 (Genschik et al., 1998). Again, we detected virtually no YFP-positive puncta in the control double phospho-mimetic variant (TTM1^S10D/S490D^, Figure 6A, B). However, MG132 treatment resulted in a significant increase in YFP-positive puncta (Figure 6C, D), suggesting that phosphorylation at the S10 and S490 residues regulates TTM1 degradation via the 26S proteasome.

**Figure 6.**
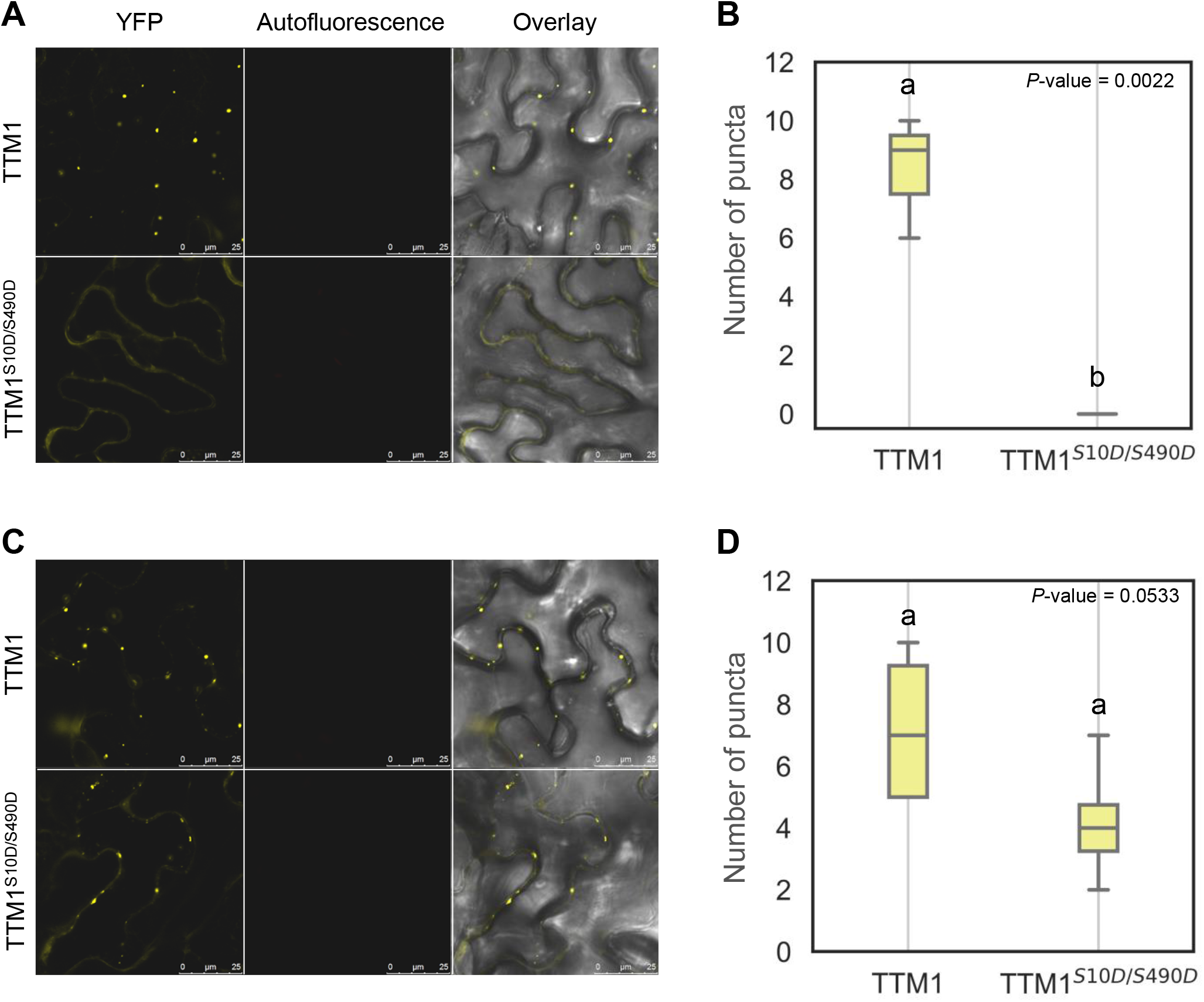
Phosphorylation of TTM1 S10 and S490 leads to protein degradation through the 26S proteasome. YFP-tagged TTM1 variants were transiently overexpressed in *N. benthamiana* leaves and visualized by confocal microscopy at 48 hpi. Leaves expressing TTM1 and phospho-mimetic TTM1^S10D/S490D^ were infiltrated at 24 hpi with **(A**,**B)** de-ionized water (control) or **(C**,**D)** 100 μM MG132. **(A, C)** Left panel: YFP signal, Middle panel: Auto fluorescence, Right panel: Overlay. **(B, D)** The number of puncta were counted from multiple confocal images and plotted as box plots displaying the first and third quartiles, split by the median; whiskers extend to a maximum of 1.5× interquartile range beyond the box. Number of images used to count puncta in (B) n=3 and (D) n>=4.

## DISCUSSION

In animals, the role of mitochondria in PCD is well established (Peña-Blanco et al., 2018). For example, the pro-apoptotic B-cell lymphoma 2 (Bcl-2) homologous antagonist/killer (BAK) and Bcl-2-associated X (BAX) proteins are tail-anchored proteins that, like TTM1, are targeted to the mitochondrial outer membrane, where they play important roles in apoptosis and tumorigenesis (Wilfling et al., 2012). Interestingly, even though no BAK and BAX homologues have been identified in plants, there is mounting evidence for a role of mitochondria in PCD in plants, mostly via a connection to reactive oxygen species (ROS) production (Wang et al., 2017; Zhang et al., 2017; Van Aken and Pogson, 2017). Recently several Arabidopsis mitochondrial outer membrane proteins (e.g. OM66 and OM47) have been connected to cell death and senescence (Zhang et al., 2014; Li et al., 2016). However, many questions regarding the role of mitochondria in plant PCD remain unanswered. In this study, we show that the mitochondrial outer membrane protein TTM1 is regulated by the plant hormone ABA at the post-translational level to accelerate senescence-related cell death, revealing a link between ABA, senescence, and mitochondria.

We had previously shown that TTM1 acts as a positive regulator of natural and dark-induced senescence (Ung and Karia et al., 2017). Here, we demonstrate that TTM1 is specifically connected to ABA-mediated senescence. Treatment with exogenous ABA, or prolonged dark treatment, lead to increased phosphorylation of TTM1 at S437 (Kline et al., 2010; Wang et al., 2013; Supplemental Table 1). Since our functional complementation analysis revealed that phosphorylation at this site is essential for TTM1 function, we embarked on the identification of the upstream signaling components (i.e. kinases) that modify TTM1 to understand the regulation of this mitochondrial protein.

In the phosphoproteomics study conducted by Wang et al., (2013), TTM1 was reported to be one of 58 proteins with increased phosphorylation upon ABA treatment and reduced phosphorylation in the *snrk2*.*2/2*.*3/2*.*6* triple mutant, suggesting that these SnRK2 kinases may directly phosphorylate TTM1 (Wang et al., 2013). This notion was supported by the delayed ABA-induced senescence phenotype in *snrk2* triple mutants, reminiscent of the *ttm1* phenotype (Gao et al., 2016; Zhao et al., 2016). However, we discovered that none of these SnRK2 kinases directly phosphorylate TTM1 (Supplemental figure S3A).

Since these SnRK2 kinases have been shown to activate at least one MAP kinase cascade upon ABA treatment, and based on the observation that the S437 residue is located within a conserved MAP kinase phosphorylation site, we focused on MAP kinases related to ABA and/or senescence. We tested ten such MAP kinases in this study, which revealed that MPK3 and MPK4, but not MPK1, MPK6, or MPK7, phosphorylate S437. MPK3 and MPK4 were not our prime candidates, since they had not previously been strongly connected to senescence. The only connection between MPK4 and senescence comes from several studies that reported ABA-mediated activation of MPK4 (Xing et al., 2008; Ichimura et al., 2000) and MKK1, the MAPKK upstream of MPK4 (Brock et al., 2010; Xing et al., 2008). Similarly, several studies reported a weak activation of MPK3 by ABA (de Zelicourt et al., 2016; Brock et al., 2010) and both kinases are transcriptionally up-regulated in senescing leaves (Supplemental Figure S4; Klepikova et al., 2016). Thus, in this study, we discovered a previously unrecognized mechanistic connection between MPK3 and MPK4 and senescence, via phosphorylation of the mitochondrial protein TTM1. Whether SnRK2 kinases also activate the upstream cascades of these MAP kinases remains to be investigated, but this may help explain the loss of TTM1 phosphorylation in the *snrk2*.*2/2*.*3/2*.*6* triple mutant (Wang et al., 2013).

In addition to the S437 residue, we identified two additional phosphorylation sites through our phospho-proteomic analyses: S10 and S490. Our confocal microscopic analysis revealed that TTM1 undergoes an intriguing change in its sub-cellular localization upon phosphorylation at these sites, and also provided evidence of TTM1 degradation. Phosphorylation is often accompanied by other post-translational modifications such as ubiquitination and interdependency between different post-translational modifications is seen for timed execution of protein function or turnover (Hunter, 2007). Previous studies have indicated that signals like light and ABA lead to protein phosphorylation, which in turn targets proteins for degradation (Cheng et al., 2017; Yang et al., 2017). The exact ubiquitination sites and the identity of the ligases that target TTM1 for degradation are under investigation. Interestingly, the animal apoptotic protein Bcl2, which also localizes at the mitochondrial outer membrane, undergoes phosphorylation at multiple sites for proper function, as well as ubiquitination for its turnover (Breitschopf et al., 2000). It will be interesting to understand the precise molecular mechanisms underlying TTM1 function and turnover in plant PCD and explore the extent to which it parallels Bcl2 in animals.

Our mass spectrometry analysis identified an additional phosphorylation event at the S535 residue upon dark treatment (Supplemental Table 1). This site does not have a typical MAP kinase target motif (Sörensson et al., 2012; Rayapuram et al., 2018) and was not phosphorylated by MPK3, MPK4, or MPK7 in our *in vitro* assays (Table 1). Therefore, TTM1 is likely regulated by another undetermined kinase at the S535 residue, suggesting its regulation by additional phosphorylation events.

TTM1 is anchored in the mitochondrial outer membrane by its C-terminal TM domain, which is followed by a tail-region of about 200 amino acids that includes a coiled-coil domain. The N-terminus of TTM1 comprises the two putative catalytic domains: the TTM and the uridine kinase-like domain. The S437 phosphorylation site is located 30 amino acids after the catalytic TTM domain, while S10 is located within the N-terminus, before the P-loop kinase domain and S490 lies just before the coiled-coil domain (Figure 3D; Supplemental Figure S5). The location of S437 indicates that phosphorylation at this position is unlikely to directly affect the enzymatic activity, since it is not in the catalytic domains, but may instead cause a general conformational change of the TTM1 protein. S437 must be an essential site for TTM1 function, as it is well conserved among both dicot and monocot TTM1 homologues. In fact, ABA treatment also leads to the phosphorylation of the same conserved site in the rice (*Oryza sativa*) TTM1 homologue (Qiu et al., 2017), further stressing the importance of phosphorylation at this residue for TTM1 function.

Taking these results together, we propose a model for TTM1 function and regulation, shown in Figure 7. ABA and other potential senescence cues lead to the activation of multiple MAP kinase cascades, resulting in the phosphorylation of TTM1 at S437 (via MPK3 and MPK4). The conformational change induced by S437 phosphorylation activates its enzymatic activity by allowing its substrate access. Thus, phosphorylation at S437 is essential for TTM1 function. This conformational change may also allow the S10 and S490 residues to be phosphorylated by multiple MAP kinases, leading to TTM1 degradation to prevent over-activation of cell death.

**Figure 7.**
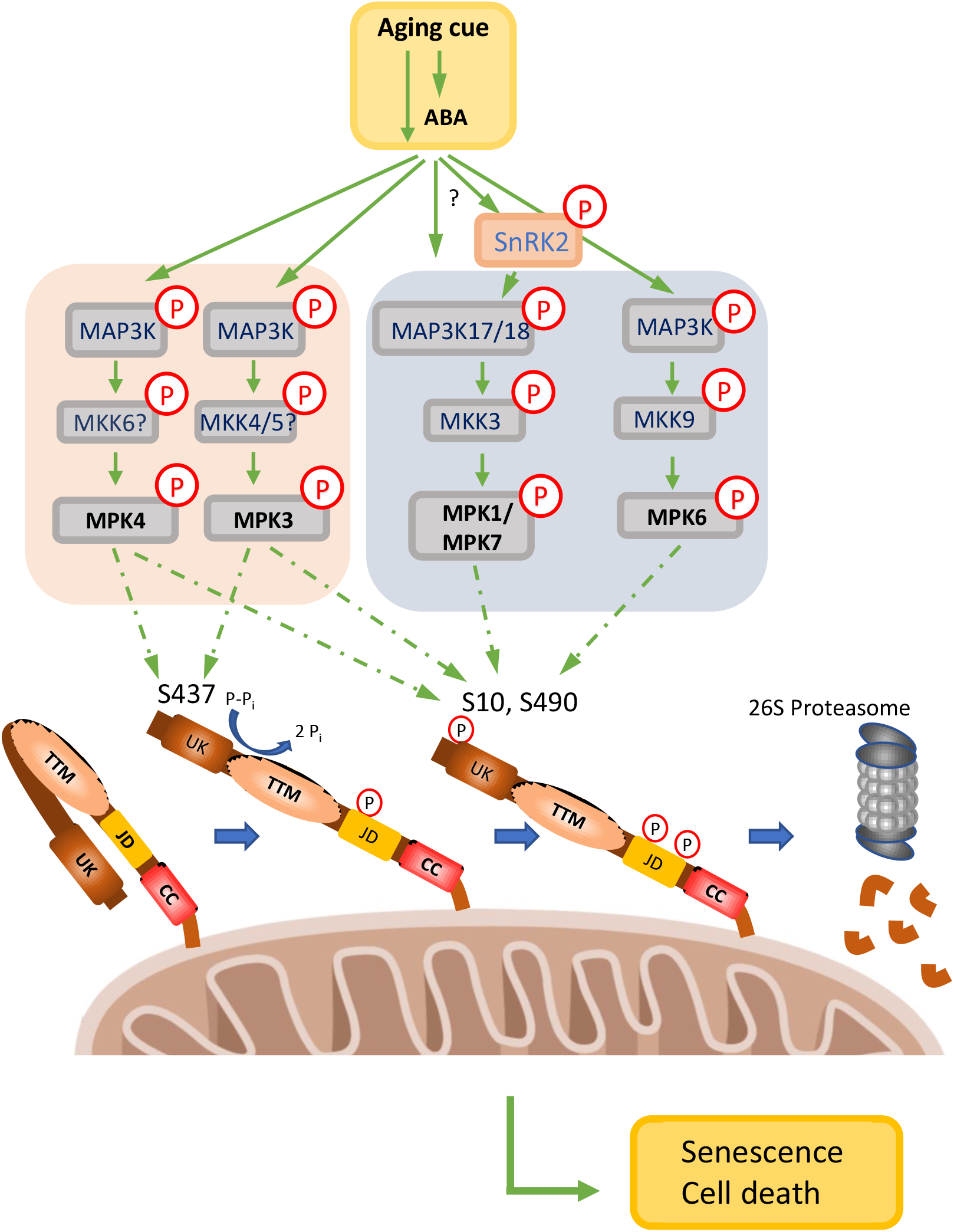
Proposed model of TTM1 function and turnover. Aging cues (or ABA) trigger the MAP kinase cascades that activate MPK3 and MPK4. They phosphorylate S437, which may cause a conformational change that leads to the activation of TTM1 (shown here as the conversion of pyrophosphate to two phosphates). This conformational change may also allow other MAP kinases (MPK6, MPK1, and MPK7) to phosphorylate S10 and S490. Once TTM1 is phosphorylated at these sites, it is marked for protein turnover by the 26S proteasome. Arrows with broken lines represent experimental findings from this study and solid lines indicate previously published data. UK = uridine kinase/P-loop kinase domain, TTM = triphosphate tunnel metalloenzyme domain, JD = junction domain, CC = Coiled-coil domain,

Our data strongly connect TTM1 to ABA signaling, which appears to regulate both the activation and the degradation of TTM1 via phosphorylation. The fact that S10/S490 phosphorylation triggers TTM1 protein turnover suggests that TTM1 is required during a specific and short window during senescence to positively regulate cell death execution. Indeed, constitutive over-expression (for instance, when driven by the 35S promoter) triggers cell death (Ung and Karia et al., 2017), suggesting that TTM1 levels have to be maintained at low levels to avoid unwanted cell death. It will be important to confirm whether the *in vivo* substrate is also pyrophosphate (Ung and Karia et al., 2017) and to establish the connection between TTM1 and other signaling components in the mitochondria.

In summary, this study reveals a previously unknown molecular link between the mitochondrial outer membrane-localized TTM1 protein and ABA signaling in senescence-related cell death. Further studies to identify the substrate of TTM1 and to understand the purpose of TTM1 localization to the mitochondrial outer membrane and its connection to PCD are in progress.

## MATERIAL AND METHODS

### Plant material and growth conditions

The *ttm1* knockout plants used in this study have been previously described (Ung et al., 2017). Arabidopsis (*Arabidopsis thaliana*) and *Nicotiana benthamiana* plants were grown on Sunshine mix in a growth chamber at 22°C, 60% relative humidity, 140 μE m^-2^ s^-1^ with a 9 h photoperiod (9 h light: 15 h dark).

### Dark and phytohormone-induced senescence assay

A combination of non-senescing leaves (leaves 4, 5, and 6, with leaf #1 being the first true leaf) of four-to five-week-old plants were detached and floated on water in petri dishes in the dark for the specified period. For phytohormone assays, leaves were floated on water containing either 50 µM ABA, 50 µM ACC, or 25 µM MeJA in a 16 h photoperiod (16 h light: 8 h dark) in a growth chamber.

### Chlorophyll quantification

Frozen leaves were ground to a fine powder before being resuspended in 80% acetone (v/v), 25 mM HEPES, pH 7.5. Total chlorophyll *a* and *b* amounts were quantified by measuring their absorbance using a spectrophotometer. We used the equation for total chlorophyll = 17.76 (*A*_646_) + 7.34 (*A*_663_), followed by normalization to fresh weight (Porra et al., 1989).

### Transgenic complementation analysis

*TTM1* variants were cloned into the pORE R2 vector, containing the *TTM1* promoter (Coutu et al.,2007; Ung et al., 2017). Stable transformation of *ttm1* plants was carried out using Agrobacterium (*Agrobacterium tumefaciens*)-mediated transformation by the floral dip method (Clough and Bent, 1998). Primer sequences are provided in Supplemental Table 2.

**Table 2:**
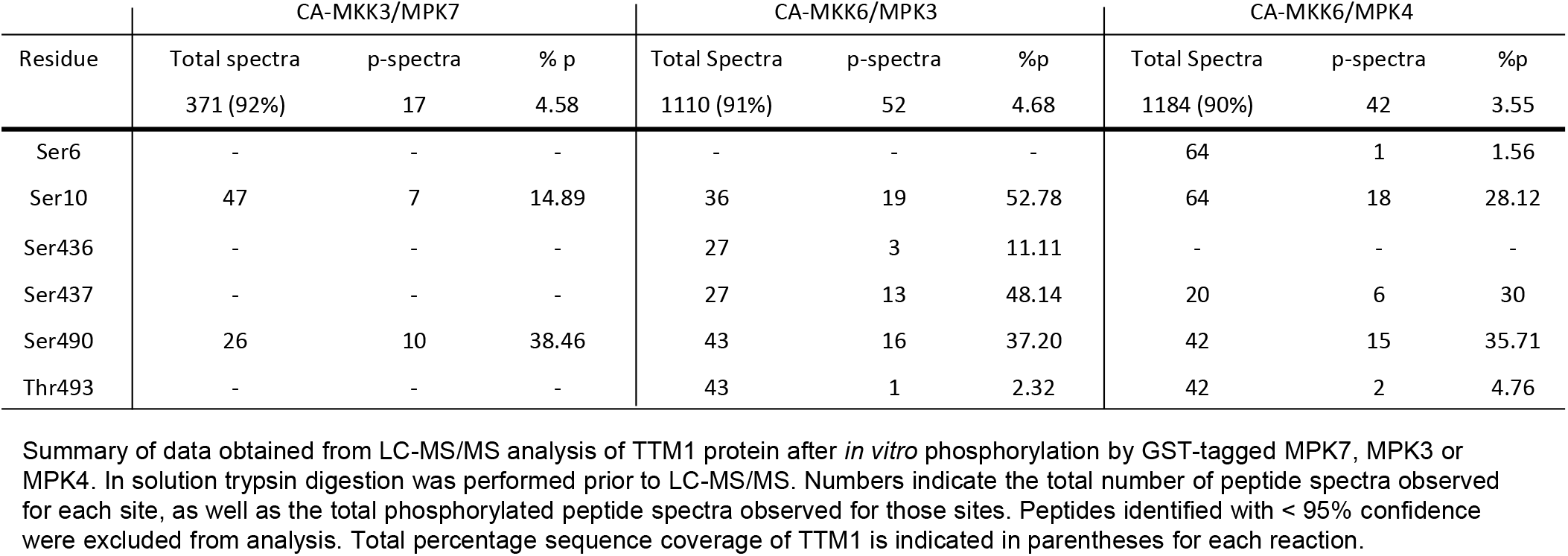
Mass spectrometry of MBP-His-TTM1 upon *in vitro* phosphorylation with MPK7, MPK3, and MPK4.

### RNA extraction, cDNA synthesis, and quantitative real time PCR

Arabidopsis leaf samples were frozen in liquid nitrogen and ground to a fine powder. RNA extraction was performed using TRIzol reagent (Life Technologies), according to the manufacturer’s instructions. First strand cDNAs were synthesized using SuperScript II Reverse Transcriptase (Life Technologies) according to the manufacturer’s instructions. RT-qPCR assays were performed using SYBR Green Master Mix (Life Technologies). Gene expression was normalized to the expression of *ELONGATION FACTOR1*-*ALPHA* (*EF1A*). Primer sequences are provided in Supplemental Table 2.

### Transient expression in N. benthamiana

Transient infiltration was performed via infiltration of 4-to 5-week-old *N. benthamiana* leaves with Agrobacterium strain GV2260. Agrobacterium cultures for all relevant constructs were grown for 6 h in Luria-Bertani (LB) broth, 10 mM MES (pH 5.7), 20 μM acetosyringone at 28ଌ, then pelleted by centrifugation and resuspended in infiltration buffer (10 mM MES, 10 mM MgCl2, 150 μM acetosyringone) to a final OD_600_ of 0.5. Agrobacterium cell suspensions carrying *CaMV35S::HC-Pro* from *Tobacco etch virus* were infiltrated alone (TEV) or co-infiltrated in a 1:1 ratio with the construct of interest. N terminally YFP-tagged TTM1 variants were cloned into pEARLEYGATE104 (Earley et al., 2006). We moved *N. benthamiana* plants to the dark 24 h after infiltration, for the indicated number of days (Ung et al., 2017).

### Bacterial induction and protein expression

TTM1 variants were cloned as MBP-His-TTM fusions in the pET28MAL vector. These constructs lacked the C-terminal TM-domain (ending at S621) (Ung et al., 2017). All kinases were GST-tagged using the pGEX-4T-3 vector. Plasmids were transformed into *Escherichia coli*, strain BL21 RIPL, for protein expression. *E. coli* containing the plasmids were grown at 37ଌ until they reached a cell density equivalent to OD_600_ = 0.4. Protein production was then induced by the addition of 0.1 mM isopropyl β-D-1-thiogalactopyranoside (IPTG); growth was allowed to continue overnight at 18ଌ before harvesting the cells by centrifugation.

### Protein purification

Bacterial pellets were resuspended in 1× phosphate buffered saline (PBS), 1% Triton X-100, 1 mM DTT, 0.5 mM PMSF, and protease inhibitor cocktail (BioShop) (GST-fusion proteins) or in 50 mM Tris-Cl, pH 7.5, 500 mM NaCl, 5% glycerol, 3 mM imidazole, 0.5 mM PMSF, and protease inhibitor cocktail (BioShop) (MBP-His fusion proteins). Cells were lysed under a French press. Soluble fractions were obtained by centrifugation at 13,000 rpm for 20 min at 4ଌ; fusion proteins were purified using either glutathione agarose (GST) or Ni-NTA agarose (MBP-His). GST proteins were eluted with 20 mM reduced glutathione, while MBP-His proteins were eluted with 200 mM imidazole. Eluted proteins were concentrated using an Amicon Ultra Centrifugal Filter Unit (50 kDa cut off), according to the manufacturer’s instructions. Protein concentration was determined by the Bradford assay at OD_595_. Proteins were stored at –80ଌ until use.

### In vitro kinase assay

For MAP kinase assays, 3 μg of MBP-His-TTM1 variants was incubated with 1 μg constitutively active (CA) GST-MKK and 1 μg GST-MPK in 25 mM Tris-Cl, pH 7.5, 10 mM MgCl_2_, 1 mM Na_3_VO_4_, 2.5 mM β-glycerophosphate, 1 mM DTT. Reactions were initiated by the addition of ATP (final concentration 100 μM ATP+2 μCi [γ-^32^P]-ATP) (Cao et al., 2018).

For SnRK2 kinase assays, 3 μg of MBP-TTM1 protein was incubated with 3 μg GST-SnRK2 in 25 mM Tris-Cl, 10 mM MgCl_2_, 1 mM DTT. Reactions were initiated by the addition of ATP (final concentration 50 μM ATP+2 μCi [γ-^32^P]-ATP) (Wang et al., 2013).

Following incubation at 30ଌ for 2 h, reactions were stopped by the addition of 1x SDS loading buffer and heating at 95ଌ for 10 min. Proteins were separated on 12% SDS-PAGE gels. Radiolabeled proteins were detected by film exposure.

### Immunoprecipitation

*N. benthamiana* plants transiently expressing *35Sp:YFP-TTM1* were used for immunoprecipitation and liquid chromatography tandem mass spectrometry ((LC-MS/MS) phosphoproteomics analysis. After 5 d of dark treatment, we collected 2 g of leaves expressing *YFP-TTM1* from control and dark samples. Tissue was frozen in liquid nitrogen and ground to fine powder. Total proteins were extracted in 50 mM Tris-HCl, pH 7.5, 50 mM NaCl, 5% glycerol, 1 mM EDTA, 5 mM DTT, 0.5% PVP, protease inhibitor cocktail (BioShop), 1% (v/v) IGEPAL, 1 mM Na_2_MoO_4_, 1 mM NaF, 0.5 mM Na_3_VO_4_; homogenized at 4ଌ for 30 min; and centrifuged at 4ଌ and 13,000 rpm for 20 min. After passing the supernatant through one layer of Miracloth, samples were diluted 1:1 with detergent-free extraction buffer. We added 20 μL of GFP-Trap (Chromotek) magnetic beads to each sample and incubated the mixtures at 4ଌ for 3 h. After centrifugation at 500*g* for 30 sec at 4ଌ. beads were washed 3 times with 1 mL 1x PBS and transferred to LoBind protein microfuge tubes. (Schwessinger et al., 2011).

### Mass spectrometry

*In vitro* kinase assays were performed as indicated above, using 5 μg MBP-His-TTM1 and 2 μg GST-MPK3/4/7 and incubated for 3 h. Kinase reactions were sent for in-solution trypsin digestion and LC-MS/MS phosphopeptide analysis at the Mass Spectrometry Facility, SPARC BioCentre (The Hospital for Sick Children, Toronto). Phosphopeptide analysis was also performed using immunoprecipitated and trypsin digested TTM1 samples from *N. benthamiana* leaves.

### Confocal microscopy

Confocal microscopy images of *N. benthamiana* leaves were collected with a Leica TCS SP5 confocal system (Leica Microsystems). Leaf discs from Agrobacterium-infiltrated areas were imaged 48 h post infiltration. The indicated samples were treated with 100 μM MG132 24 h prior to visualization. YFP (520-590 nm) or chloroplast autofluorescence (650-700 nm) was detected under a 63x oil immersion objective lens with 3x zoom, using the 514-nm OPSL laser set to 33%.

### Statistical analysis

Statistical analysis was performed using one-way analysis of variance (ANOVA) and Tukey’s post hoc test using Python scripts in Jupyter notebook and Student’s *t*-test using Microsoft Excel.

## Accession numbers

Sequence data from this article can be found in the Arabidopsis Genome Initiative or GenBank/EMBL databases under the following accession numbers: *Arabidopsis thaliana TTM1* (At1g73980), *EF1A* (At5g60390), *β-tub* (<u>At5g23860</u>), *MPK1* (At1g10210), *MPK2 (*At1g59580*), MPK7* (At2g18170), *MPK14* (At4g36450), *MPK6* (At2g43790), *MPK3* (At3g45640), *MPK4* (At4g01370), *MPK11* (At1g01560), *SnRK2*.*2* (At3g50500), *SnRK2*.3 (At5g66880), *SnRK2*.*6* (At4g33950), *RD29A* (At5g52310), *SAG12* (At5g45890)

## Acknowledgements

This work was supported by the Canadian Foundation for Innovation and the Ontario Research Fund to KY and an Ontario graduate Scholarship to PK. We thank Dr. Brian Ellis and Dr Jin Suk Lee for providing us MAP kinase clones. We also thank Dr. Thomas DeFalco for providing expertise in protein techniques.

## Author contributions

W.M., P.K. and K.Y. designed the research, P.K., and W.M. performed the research, P.K., W.M. and K.Y. analyzed the data, W.M. and P.K. wrote the article.

## REFERENCES

Bettendorff, L., and Wins, P. (2013) Thiamine triphosphatase and the CYTH superfamily of proteins. FEBS J. 24: 6443–6455.

Breeze, E., Harrison, E., McHattie, S., Hughes, L., Hickman, R., Hill, C., Kiddle, S., Kim, Y.S., Penfold, C.A., Jenkins D et al. (2011) High-resolution temporal profiling of transcripts during Arabidopsis leaf senescence reveals a distinct chronology of processes and regulation. Plant Cell 23: 873–894.

Breitschopf, K., Haendeler, J., Malchow, P., Zeiher, A.M., Dimmeler, S. (2000) Posttranslational Modification of Bcl-2 Facilitates Its Proteasome-Dependent Degradation: Molecular Characterization of the Involved Signaling Pathway. Mol. Cell Biol. 20: 1886–1896.

Brock, A.K., Willmann, R., Kolb, D., Grefen, L., Lajunen, H.M., Bethke, G., Lee, J., Nürnberger, T., Gust, A.A. (2010) The Arabidopsis mitogen-activated protein kinase phosphatase PP2C5 affects seed germination, stomatal aperture, and abscisic acid-inducible gene expression. Plant Physiol. 153: 1098–1111.

Cao, F.Y., DeFalco, T.A., Moeder, W., Li, B., Gong, Y., Liu, X.M., Taniguchi, M., Lumba, S., Toh, S., Shan, L., Ellis, B., Desveaux, D., Yoshioka, K. (2018) Arabidopsis ETHYLENE RESPONSE FACTOR 8 (ERF8) has dual functions in ABA signaling and immunity. BMC Plant Biology 18: 211.

Cheng, C., Wang, Z., Ren, Z., Zhi, L., Yao, B., Su, C., Liu, L., and Li, X. (2017). SCFAtPP2-B11modulates ABA signaling by facilitating SnRK2.3 degradation in Arabidopsis thaliana. PLoS Genet. 13: e1006947.

Chrobok, D., Law, S.R., Brouwer, B., Lindén, P., Ziolkowska, A., Liebsch, D., Narsai, R., Szal, B., Moritz, T., Rouhier, N., Whelan, J., Gardeström, P., and Keech, O. (2016) Dissecting the Metabolic Role of Mitochondria during Developmental Leaf Senescence. Plant Physiol. 172: 2132–2153.

Clough, S.J. and Bent, A.F. (1998). Floral dip: A simplified method for Agrobacterium-mediated transformation of Arabidopsis thaliana. Plant J. 16: 735–743.

Coutu, C., Brandle, J., Brown, D., Brown, K., Miki, B., Simmonds, J., and Hegedus, D. (2007) pORE: a modular binary vector series suited for both monocot and dicot plant transformation. Transgenic Res. 16: 771–781.

Cutler, S.R., Rodriguez, P.L., Finkelstein, R.R., and Abrams, S.R. (2010) Abscisic acid: emergence of a core signaling network. Annu. Rev. Plant Biol. 61:651–679.

Daneva, A., Gao, Z., Van Durme, M., Nowack, M.K. (2016) Functions and Regulation of Programmed Cell Death in Plant Development. Annu. Rev. Cell Dev. Biol. 6: 441–468.

Danquah, A., de Zélicourt, A., Boudsocq, M., Neubauer, J., Frei Dit Frey, N., Leonhardt, N., Pateyron, S., Gwinner, F., Tamby, J.P., Ortiz-Masia, D., Marcote, M.J., Hirt, H., and Colcombet, J. (2015). Identification and characterization of an ABA-activated MAP kinase cascade in Arabidopsis thaliana. Plant J. 82: 232–244.

de Zelicourt, A., Colcombet, J., and Hirt, H. (2016) The Role of MAPK Modules and ABA during Abiotic Stress Signaling. Trends Plant Sci. 21: 677–685.

Earley, K., Haag, J.R., Pontes, O., Opper, K., Juehne, T., Song, K., Pikaard, C.S. (2006) Gateway-compatible vectors for plant functional genomics and proteomics. Plant J. 45: 616–629.

Fujii, H., and Zhu, J.K. (2009). Arabidopsis mutant deficient in 3 abscisic acid-activated protein kinases reveals critical roles in growth, reproduction, and stress. Proc. Natl. Acad. Sci. USA 106: 8380–8385.

Fujita, Y., Nakashima, K., Yoshida, T., Katagiri, T., Kidokoro, S., Kanamori, N., Umezawa, T., Fujita, M., Maruyama, K., Ishiyama, K., Kobayashi, M., Nakasone, S., Yamada, K., Ito, T., Shinozaki, K., Yamaguchi-Shinozaki, K. (2009) Three SnRK2 protein kinases are the main positive regulators of abscisic acid signaling in response to water stress in Arabidopsis. Plant Cell Physiol. 50: 2123–2132.

Gallagher, D.T., Smith, N.N., Kim, S.K., Heroux, A., Robinson, H., Reddy, P.T. (2006) Structure of the class IV adenylyl cyclase reveals a novel fold. J. Mol. Biol. 362: 114–122.

Gao, S., Gao, J., Zhu, X., Song, Y., Li, Z., Ren, G., Zhou, X., Kuai, B. (2016) ABF2, ABF3, and ABF4 Promote ABA-Mediated Chlorophyll Degradation and Leaf Senescence by Transcriptional Activation of Chlorophyll Catabolic Genes and Senescence-Associated Genes in Arabidopsis. Mol Plant. 9: 1272–1285.

Genschik, P., Criqui, M.C., Parmentier, Y., Derevier, A., and Fleck, J. (1998). Cell cycle-dependent proteolysis in plants: Identification of the destruction box pathway and metaphase arrest produced by the proteasome inhibitor MG132. Plant Cell 10: 2063–2075.

Goda H, Sasaki E, Akiyama K, Maruyama-Nakashita A, Nakabayashi K, Li W, Ogawa M, Yamauchi Y, Preston J, Aoki K, Kiba T, Takatsuto S, Fujioka S, Asami T, Nakano T, Kato H, Mizuno T, Sakakibara H, Yamaguchi S, Nambara E, Kamiya Y, Takahashi H, Hirai MY, Sakurai T, Shinozaki K, Saito K, Yoshida S, Shimada Y. (2008) The AtGenExpress hormone and chemical treatment data set: experimental design, data evaluation, model data analysis and data access. Plant J. 55: 526–542.

Heese, A., Hann, D.R., Gimenez-Ibanez, S., Jones, A.M.E., He, K., Li, J., Schroeder, J.I., Peck, S.C., and Rathjen, J.P. (2007). The receptor-like kinase SERK3/BAK1 is a central regulator of innate immunity in plants. Proc. Natl. Acad. Sci. U. S. A. 104: 12217–12222.

Hunter, T. (2007). The Age of Crosstalk: Phosphorylation, Ubiquitination, and Beyond. Mol. Cell 28: 730–738.

Ichimura, K., Mizoguchi, T., Yoshida, R., Yuasa, T., Shinozaki, K. (2000) Various abiotic stresses rapidly activate Arabidopsis MAP kinases ATMPK4 and ATMPK6. Plant J. 24: 655–665.

Iyer, L.M. and Aravind, L. (2002) The catalytic domains of thiamine triphosphatase and CyaB-like adenylyl cyclase define a novel superfamily of domains that bind organic phosphates. BMC Genomics 3: 33.

Jammes, F., Song, C., Shin, D., Munemasa, S., Takeda, K., Gu, D., Cho, D., Lee, S., Giordo, R., Sritubtim, S., Leonhardt, N., Ellis, B.E., Murata, Y., Kwak, J.M. (2009) MAP kinases MPK9 and MPK12 are preferentially expressed in guard cells and positively regulate ROS-mediated ABA signaling. Proc. Natl. Acad. Sci. U. S. A. 106: 20520–20525.

Keech, O., Pesquet, E., Ahad, A., Askne, A., Nordvall, D., Vodnala, S.M., Tuominen, H., Hurry, V., Dizengremel, P., Gardestrom, P. (2007) The different fates of mitochondria and chloroplasts during dark-induced senescence in Arabidopsis leaves. Plant Cell Environ 30: 1523–1534.

Keppetipola, N., Jain, R., and Shuman, S. (2007) Novel triphosphate phosphohydrolase activity of Clostridium thermocellum TTM, a member of the triphosphate tunnel metalloenzyme superfamily. J Biol. Chem. 282: 11941–11949.

Kim, J., Kim, J.H., Lyu, J.I., Woo, H.R., and Lim, P.O. (2018) New insights into the regulation of leaf senescence in Arabidopsis. J. Exp. Bot. 69: 787–799.

Kim, J.H Chung, K.M Woo, H.R. (2011) Three positive regulators of leaf senescence in Arabidopsis, ORE1, ORE3 and ORE9, play roles in crosstalk among multiple hormone-mediated senescence pathways. Genes Genomics 33: 373–381.

Klepikova, A.V., Kasianov, A.S., Gerasimov, E.S., Logacheva, M.D., Penin, A.A. (2016) A high resolution map of the Arabidopsis thaliana developmental transcriptome based on RNA-seq profiling. Plant J. 88:1058–1070.

Kline, K.G., Barrett-Wilt, G.A., and Sussman, M.R. (2010) In planta changes in protein phosphorylation induced by the plant hormone abscisic acid. Proc. Natl. Acad. Sci. U. S. A. 107: 15986–15991.

Kmiecik, P., Leonardelli, M., and Teige, M. (2016) Novel connections in plant organellar signalling link different stress responses and signalling pathways. J. Exp. Bot. 67:3793–3807.

Kriechbaumer. V., Shaw, R., Mukherjee, J., Bowsher, C.G., Harrison, A.M., and Abell, B.M. (2009). Subcellular distribution of tail-anchored proteins in Arabidopsis. Traffic 10: 1753–1764.

Lakaye, B., Makarchikov, A.F., Wins, P., Margineanu, I., Roland, S., Lins, L., Aichour, R., Lebeau, L., El Moualij, B., Zorzi, W., Coumans, B., Grisar, T., Bettendorff, L. (2004) Human recombinant thiamine triphosphatase: purification, secondary structure and catalytic properties. International J. Biochem. Cell Biol. 36: 1348–1364.

Li, L., Kubiszewski-Jakubiak, S., Radomiljac, J., Wang, Y., Law, S.R., Keech, O., Narsai, R., Berkowitz, O., Duncan, O., Murcha, M.W., Whelan, J. (2016) Characterization of a novel β-barrel protein (AtOM47) from the mitochondrial outer membrane of Arabidopsis thaliana. J. Exp. Bot. 67: 6061–6075.

Lima, C.D., Wang, L.K., and Shuman, S. (1999) Structure and mechanism of yeast RNA triphosphatase: an essential component of the mRNA capping apparatus. Cell 99: 533–543.

Liu, T., Longhurst, A.D., Talavera-Rauh, F., Hokin, S.A., Barton, M.K. (2016) The Arabidopsis transcription factor ABIG1 relays ABA signaled growth inhibition and drought induced senescence. eLife 5: e13768.

Lorenzo-Orts, L., Hohmann, U., Zhy, J., Horthorn, M. (2019) Molecular characterization of Chad domains as inorganic polyphosphate-binding modules. Life Sci. Alliance 2: 1–14.

Martinez, J., Truffault, V., and Hothorn, M. (2015). Structural Determinants for Substrate Binding and Catalysis in Triphosphate Tunnel Metalloenzymes. J. Biol. Chem. 290: 23348–23360.

Matsuoka, D., Yasufuku, T., Furuya, T., Nanmori, T. (2015) An abscisic acid inducible Arabidopsis MAPKKK, MAPKKK18 regulates leaf senescence via its kinase activity. Plant Mol. Biol. 87: 565–575.

Moeder, W., Garcia-Petit, C., Ung, H., Fucile, G., Samuel, M.A., Christendat, D., Yoshioka, K. (2013) Crystal structure and biochemical analyses reveal that the Arabidopsis Triphosphate Tunnel Metalloenzyme, AtTTM3, is a tripolyphosphatase and is involved in root development. Plant J. 76: 615–626.

Peña-Blanco, A., García-Sáez, A.J. (2018) Bax, Bak and beyond - mitochondrial performance in apoptosis. FEBS J. 285: 416–431.

Porra, R.J., Thompson, W.A., Kriedemann, P.E. (1989) Determination of accurate extinction coefficients and simultaneous equations for assaying chlorophylls a and b extracted with four different solvents: verification of the concentration of chlorophyll standards by atomic absorption spectroscopy. Biochim. Biophys. Acta 975: 384–394.

Qiu, J., Hou, Y., Wang, Y., Li, Z., Zhao, J., Tong, X., Lin, H., Wei, X., Ao, H., Zhang, J. (2017) A Comprehensive Proteomic Survey of ABA-Induced Protein Phosphorylation in Rice (Oryza sativa L.). Int. J. Mol. Sci. 3;18(1).

Rayapuram, N., Bigeard, J., Alhoraibi, H., Bonhomme, L., Hesse, A.M., Vinh, J., Hirt, H., Pflieger, D. (2018) Quantitative Phosphoproteomic Analysis Reveals Shared and Specific Targets of Arabidopsis Mitogen-Activated Protein Kinases (MAPKs) MPK3, MPK4, and MPK6. Mol. Cell. Proteomics 17:61–80.

Samet, J.S., and Sinclair, T.R. (1980) Leaf Senescence and Abscisic Acid in Leaves of Field-grown Soybean. Plant Physiol. 66:1164–1168.

Schwessinger, B., Roux, M., Kadota, Y., Ntoukakis, V., Sklenar, J., Jones, A., and Zipfel, C. (2011). Phosphorylation-dependent differential regulation of plant growth, cell death, and innate immunity by the regulatory receptor-like kinase BAK1. PLoS Genet. 7: e1002046.

Sörensson, C., Lenman, M., Veide-Vilg, J., Schopper, S., Ljungdahl, T., Grøtli, M., Tamás, M.J., Peck, S.C., Andreasson, E. (2012) Determination of primary sequence specificity of Arabidopsis MAPKs MPK3 and MPK6 leads to identification of new substrates. Biochem. J. 446: 271–218.

Sussmilch, F.C., Schultz, J., Hedrich, R., Roelfsema, M.R.G. (2019) Acquiring Control: The Evolution of Stomatal Signalling Pathways. Trends Plant Sci. 24:342–351.

Tajdel, M., Mituła, F., Ludwików, A. (2016) Regulation of Arabidopsis MAPKKK18 by ABI1 and SnRK2, components of the ABA signaling pathway. Plant Signal. Behav. 11: e1139277.

Umezawa, T., Sugiyama, N., Takahashi, F., Anderson, J.C., Ishihama, Y., Peck, S.C., and Shinozaki, K. (2013). Genetics and phosphoproteomics reveal a protein phosphorylation network in the abscisic acid signaling pathway in Arabidopsis thaliana. Sci. Signal. 6, rs8.

Ung, H., Karia, P., Ebine, K., Ueda, T., Yoshioka, K., Moeder, W. (2017) Triphosphate Tunnel Metalloenzyme Function in Senescence Highlights a Biological Diversification of This Protein Superfamily. Plant Physiol. 175, 473–485.

Ung, H., Moeder, W., and Yoshioka, K. (2014) Arabidopsis triphosphate tunnel metalloenzyme2 (AtTTM2) is a negative regulator of the salicylic acid-mediated feedback amplification loop for defense responses. Plant Physiol. 166: 1009–1021.

Van Aken, O., Pogson, B.J. (2017) Convergence of mitochondrial and chloroplastic ANAC017/PAP-dependent retrograde signalling pathways and suppression of programmed cell death. Cell Death Differ. 24: 955–960.

Wada, S., Ishida, H., Izumi, M., Yoshimoto, K., Ohsumi, Y., Mae, T., Makino, A. (2009) Autophagy plays a role in chloroplast degradation during senescence in individually darkened leaves. Plant Physiol. 149: 885–93.

Wang Y, Peng X, Yang Z, Zhao W, Xu W, Hao J, Wu W, Shen XL, Luo Y, Huang K. (2017) iTRAQ Mitoproteome Analysis Reveals Mechanisms of Programmed Cell Death in Arabidopsis thaliana Induced by Ochratoxin A. Toxins (Basel) 9: pii: E167.

Wang, P., Xue, L., Batelli, G., Lee, S., Hou, Y.J., Van Oosten, M.J., Zhang, H., Tao, W.A., and Zhu, J.K. (2013). Quantitative phosphoproteomics identifies SnRK2 protein kinase substrates and reveals the effectors of abscisic acid action. Proc. Natl. Acad. Sci. USA 110, 11205–11210.

Weaver, L.M., Gan, S., Quirino, B., Amasino, R.M. (1998) A comparison of the expression patterns of several senescence associated genes in response to stress and hormone treatment. Plant Mol. Biol. 37: 455–469.

Wilfling, F., Weber, A., Potthoff, S., Vögtle, F.N., Meisinger, C., Paschen, S.A., Häcker, G. (2012) BH3-only proteins are tail-anchored in the outer mitochondrial membrane and can initiate the activation of Bax. Cell Death Differ. 19: 1328–1336.

Xing, Y., Jia, W., Zhang, J. (2008) AtMKK1 mediates ABA-induced CAT1 expression and H2O2 production via AtMPK6-coupled signaling in Arabidopsis. Plant J. 54: 440–451.

Xu, J., Zhang, S. (2015) Mitogen-activated protein kinase cascades in signaling plant growth and development. Trends Plant Sci. 20:56–64.

Yan, A., Chen, Z. (2017) The pivotal role of abscisic acid signaling during transition from seed maturation to germination. Plant Cell Rep. 36:689–703.

Yang, J., Worley, E., and Udvardi, M. (2014) A NAP-AAO3 regulatory module promotes chlorophyll degradation via ABA biosynthesis in Arabidopsis leaves. Plant Cell 26, 4862–4874.

Yang, W., Zhang, W., Wang, X. (2017) Post-translational control of ABA signalling: the roles of protein phosphorylation and ubiquitination. Plant Biotechnol. J. 15: 4–14.

Zhang, B., Van Aken, O., Thatcher, L., De Clercq, I., Duncan, O., Law, S.R., Murcha, M.W., van der Merwe, M., Seifi, H.S., Carrie, C., Cazzonelli, C., Radomiljac, J., Höfte, M., Singh, K.B., Van Breusegem, F., Whelan, J. (2014) The mitochondrial outer membrane AAA ATPase AtOM66 affects cell death and pathogen resistance in Arabidopsis thaliana. Plant J. 80: 709–727.

Zhang, S., Li, C., Wang, R., Chen, Y., Shu, S., Huang, R., Zhang, D., Li, J., Xiao, S., Yao, N., Yang, C. (2017) The Arabidopsis Mitochondrial Protease FtSH4 Is Involved in Leaf Senescence via Regulation of WRKY-Dependent Salicylic Acid Accumulation and Signaling. Plant Physiol. 173: 2294–2307.

Zhao, Y., Chan, Z., Gao, J., Xing, L., Cao, M., Yu, C., Hu, Y., You, J., Shi, H., Zhu, Y., et al. (2016). ABA receptor PYL9 promotes drought resistance and leaf senescence. Proc. Natl. Acad. Sci. USA 113: 1949–1954.

Zhou, C., Cai, Z., Guo, Y., and Gan, S. (2009). An Arabidopsis mitogen activated protein kinase cascade, MKK9-MPK6, plays a role in leaf senescence. Plant Physiol. 150: 167–177.

Clough, S.J. and Bent, A.F. (1998). Floral dip: Asimplified method for Agrobacterium-mediated transformation of Arabidopsis thaliana. Plant J. 16: 735–743.

Kim, J.H· Chung, K.M· Woo, H.R. (2011) Three positive regulators of leaf senescence in Arabidopsis, ORE1, ORE3 and ORE9, play roles in crosstalk among multiple hormone-mediated senescence pathways. Genes Genomics 33: 373–381.

Klepikova, A.V., Kasianov, A.S., Gerasimov, E.S., Logacheva, M.D., Penin, A.A. (2016) Ahigh resolution map of the Arabidopsis thaliana developmental transcriptome based on RNA-seq profiling. Plant J. 88:1058–1070.

Lima, C.D., Wang, L.K., and Shuman, S. (1999) Structure and mechanism of yeast RNAtriphosphatase: an essential component of the mRNA capping apparatus. Cell 99: 533–543.

Qiu, J., Hou, Y., Wang, Y., Li, Z., Zhao, J., Tong, X., Lin, H., Wei, X., Ao, H., Zhang, J. (2017) AComprehensive Proteomic Survey of ABA-Induced Protein Phosphorylation in Rice (Oryza sativa L.). Int. J. Mol. Sci. 3;18(1).

Weaver, L.M., Gan, S., Quirino, B., Amasino, R.M. (1998) Acomparison of the expression patterns of several senescence associated genes in response to stress and hormone treatment. Plant Mol. Biol. 37: 455–469.

Yang, J., Worley, E., and Udvardi, M. (2014) ANAP-AAO3 regulatory module promotes chlorophyll degradation via ABA biosynthesis in Arabidopsis leaves. Plant Cell 26, 4862–4874.

